# Single-cell transcriptomics of adult skin VE-cadherin expressing lineages during hair cycle

**DOI:** 10.1101/2023.03.22.533784

**Authors:** Gopal Chovatiya, Kefei Nina Li, Sangeeta Ghuwalewala, Tudorita Tumbar

## Abstract

Adult skin homeostasis involves global reorganization of dermal lineages at different stages of the mouse hair growth cycle. Vascular endothelial cadherin (VE-cadherin encoded by *Cdh5*) expressing cells from blood and lymphatic vasculature structures are known to remodel during the adult hair cycle. Here we employ single-cell RNA-sequencing (scRNA-seq) 10x-genomics analysis of FACS-sorted VE-cadherin expressing cells marked via Cdh5-CreER genetic labeling at resting (telogen) and growth (anagen) stage of hair cycle. Our comparative analysis between the two stages uncovers a persistent Ki67^+^ proliferative EC population and documents changes in EC population distribution and gene expression. Global gene expression changes in all the analyzed populations revealed bioenergetic metabolic changes that may drive vascular remodeling during HF growth phase, alongside a few highly restricted cluster-specific gene expression differences. This study uncovers active cellular and molecular dynamics of adult skin endothelial lineages during hair cycle that may have broad implications in adult tissue regeneration and for understanding vascular disease.

## Introduction

Vascular endothelial (VE) cadherin (encoded by *Cdh5*) marks endothelial cell (EC), and is expressed by both blood vessels (capillaries, veins, and arteries) and lymphatic vessels^1^. The encoding gene and mRNA are referred to as *Cdh5* and the protein product as VE-cadherin, a convention we follow throughout our paper. VE-cadherin is a cell-cell contact molecule, recognized for its essential role in the formation and maturation of vascular tubes^1, 2^. It also contributes to signaling within the ECs by interacting with receptors of the VEGF, Notch, FGF, and the TGF-β pathway^3^. Its molecular interactions ensure vasculature resilience, flexibility, mobility, and permeability, thus endowing ECs with their specific characteristics^3, 4, 5, 6^. Non-endothelial tumor cells can also aberrantly activate VE-cadherin, forming tubular irrigation networks that assist tumor growth^7^. Single-cell (sc) RNA-seq analysis of ECs from multiple adult organs (but not skin) uncovered significant tissue-specific gene expressions^8, 9, 10^, but generally identified the similar types of mature ECs. Single-cell transcriptomics in the adult mouse dorsal aorta identified an immature *Cdh5^+^* cellular subset deemed an endothelial vascular progenitor (EVP)^11^, possibly also present in skin wounds and tumors^12, 13^. Little is known about the dynamic changes in VE-cadherin lineage heterogeneity and molecular makeup during adult tissue homeostasis, in the absence of injury or cancer.

Adult skin and hair follicles are an excellent model system to study the organization of adult VE-cadherin^+^ lineages during tissue homeostasis. Skin is a highly regenerative tissue that requires an ongoing supply of O_2_ and nutrients. The hair follicles, made largely of epithelial cells undergo periodic transitions from resting/quiescence (telogen) to growth (anagen) phase during the hair homeostatic cycle. These phases are synchronous in mice for the first cycle, providing an opportunity to access homeostatic processes that are generally more hidden in less synchronous adult tissues. At anagen, the skin hypodermis significantly expands in size and becomes filled with engorging adipocytes and blood vessels that wrap around the downward growing hair follicle bulbs while the lymphatic vessels also remodel and change their drainage capacity^14, 15^ (Fig. 1a). Other skin structures, such as nerves and sub-cutaneous muscles, may also be remodeled during the hair cycle^16^, but this is not well understood. Characterization of the cellular heterogeneity of skin VE-cadherin^+^ lineage organization and the changes in molecular makeup as this lineage remodels during adult skin homeostasis is currently lacking.

**Fig. 1.**
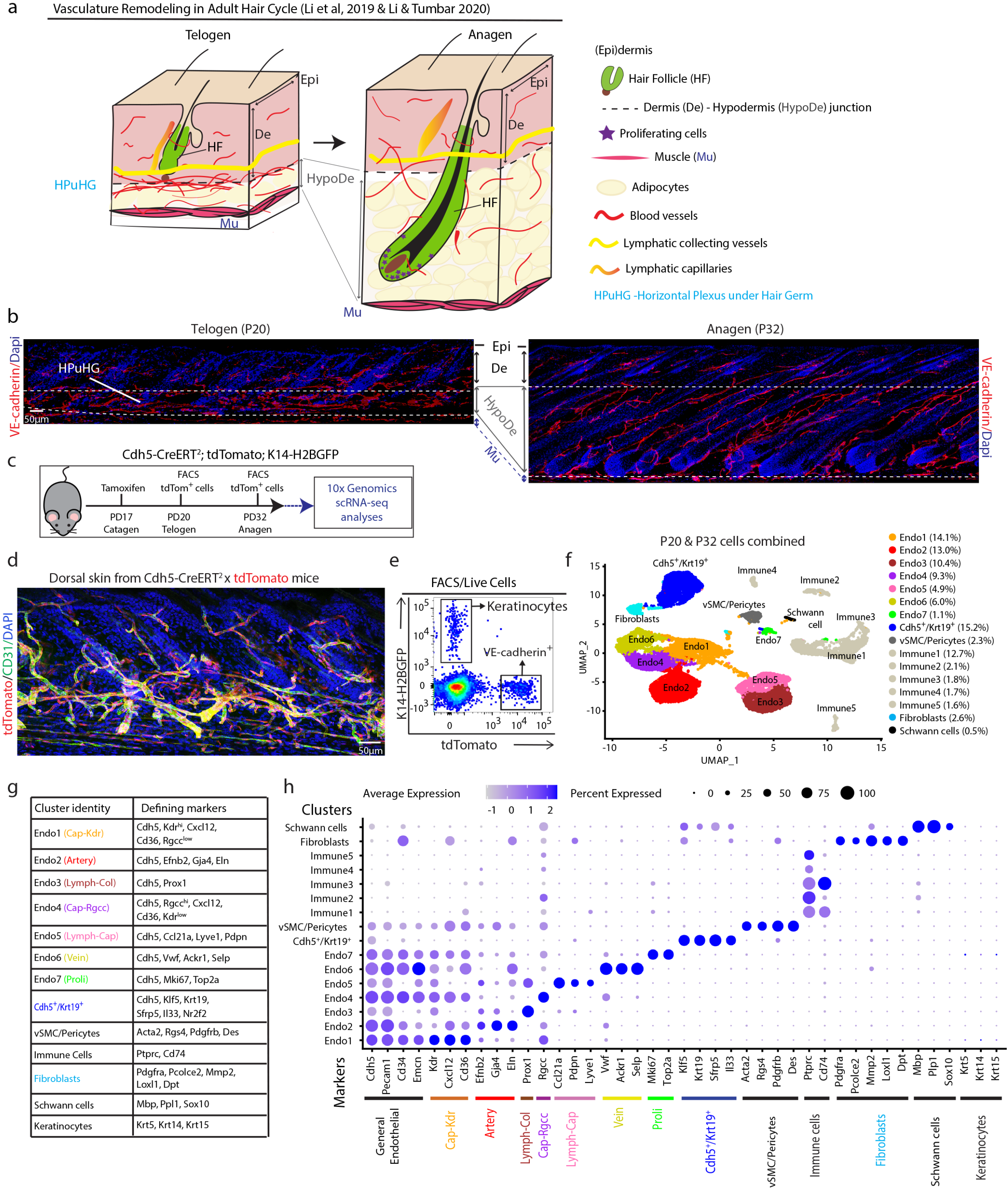
Adult skin VE-cadherin-marked cells obtained by scRNA-seq. **a** Schematic highlighting vasculature remodeling during adult mouse hair cycle (Li *et al*., 2021). Note the <u>H</u>orizontal <u>P</u>lexus <u>u</u>nderneath the <u>H</u>air <u>G</u>erm (<u>HPuHG</u>) in telogen and its dispersal with vessels wrapping around hair follicle bulbs at anagen. Lymphatic capillaries move slightly away from the hair and widen their caliber at anagen (Gur-Cohen *et al*., 2019). **b** Maximal projections of confocal optical z-stacks of 70μm thick dorsal skin sections stained with VE-cadherin (red) and Dapi (blue). The HPuHG area is marked between white dotted line. See also Supplementary Fig. 1a, b. Scale bar 50μm. Multiple adjacent images were combined using the stitching function in Fiji. **c** Schematic showing tamoxifen induction and sample collection timepoints of *Cdh5*-CreERT^2^; tdTomato; *Krt14*-H2BGFP mice for scRNA-seq analysis. **d** Immunofluorescence imaging of *Cdh5*-tdTomato (red) in dorsal skin tissue sections stained for CD31 (green), scale bar 50μm. **e** FACS purification of tdTomato^+^ cells from dorsal skin of *Cdh5*-CreERT^2^ x tdTomato; *Krt14*-H2BGFP mice, n=2 mice/stage. **f** <u>U</u>niform <u>M</u>anifold <u>A</u>pproximation and <u>P</u>rojection (<u>UMAP</u>) plot generated after combining all 4 samples (2 samples/stage) showing all 16 cell populations, their first level identity and their relative abundance in %. All four samples were integrated using the *Harmony* package before UMAP analysis. See also Supplementary Fig. 1i, j. **g** Table of previously known markers that were used for population identification. **h** Dotplot showing expression of the population identifying markers.

To characterize the heterogeneity and dynamics of mouse skin VE-cadherin^+^ lineages, we employed 10x Genomics scRNA-seq of sorted *Cdh5*-CreERT^2^;tdTomato labelled cells at two hair cycle stages: quiescence (telogen) and growth (anagen). We also document the extent of adult EC proliferation during hair cycle and uncover significant changes in blood and lymphatic EC fractions. We map the genome wide changes in gene expression that highlight the global rewiring of cellular metabolic signatures, signaling pathways, and other biological processes. This demonstrates the active dynamics of the VE-cadherin lineage in adult skin homeostasis, contrasting its more quiescent nature in other adult tissues^17, 18, 19^. Our study uncovers cellular and functional heterogeneity of the VE-cadherin expressing lineage and its active dynamics during adult tissue homeostasis. This work opens new avenues for further investigation into endothelial population specific functional roles in adult regenerative tissues with potential implications for vascular disease.

## Results

### Labeling and isolation of adult skin VE-cadherin^+^ (*Cdh5*) cells documents growth rates

Previously, we and others documented changes in skin vasculature organization during hair cycle^20, 21, 22^, as adult skin considerably expands in thickness at the transition from telogen to anagen^23^ (Fig. 1a). The hair follicle grows downward, the hypodermis fills up with fat cells derived from skin resident adipose progenitor cells, and the vasculature stretches out and nourishes tissue growth^24^. In particular, the lymphatic capillaries were shown to increase in caliber and move away from the HFs^25, 26^. We showed by quantification of skin sections that a <u>h</u>orizontal vascular <u>p</u>lexus (CD31^+^/ VE-cadherin^+^) <u>u</u>nderneath the <u>h</u>air <u>g</u>erm (HPuHG) disperses out into the hypodermis at the telogen-anagen transition^20^ (Fig. 1a). Whole mount skin imaging (Li et al, in press) and 70μm skin sections that were VE-cadherin stained or *Cdh5*-CreERT^2^ x tdTomato (*Cdh5*-tdTomato^+^) labeled, followed by 3D confocal microscopy confirmed the previously reported^20^ large-scale remodeling (Fig. 1b and Supplementary Fig. 1a, b). We and others also reported an increase in Ki67^+^ or BrdU^+^ proliferative CD31^+^ ECs at anagen^20, 21, 22^. To quantitatively evaluate the magnitude of EC proliferation in homeostasis, we performed BrdU labeling from PD17-25, encompassing the hair cycle transition from quiescence to growth. We used *Cdh5*-CreERT^2^ x tdTomato (*Cdh5*-tdTomato^+^)^27, 28^ reporter mice induced with tamoxifen (TM) at postnatal day (PD)17, followed by FACS isolation of tdTomato^+^ ECs (Supplementary Fig. 1c, d). Confocal microscopy of skin sections confirmed both high BrdU labeling efficiency and the presence of BrdU^+^/VE-cadherin^+^ vascular cells in single optical images (Supplementary Fig. 1e). Analysis of FACS sorted tdTomato^+^ cells deposited on slides and stained with antibodies for BrdU and VE-cadherin (*Cdh5*) (Supplementary Fig. 1f) showed that ~80% of the sorted tdTomato^+^ were strongly VE-cadherin-positive by immunofluorescence (IF) staining, and of those ~15% incorporated BrdU during the 8-day labeling period (Supplementary Fig. 1g). This suggests that in the period that accompanies the transition from hair quiescence to growth ~ 1-2% ECs divide on average/day. Proliferation was reported to continue through early anagen and peaks at full anagen^20, 21, 22^. Thus, we can estimate that at least a third of skin VE-cadherin expressing cells engage in proliferation during this hair cycle in mice. Combined with previous characterization^20^, these data documents substantial cellular dynamics in the VE-cadherin^+^ skin compartment during hair cycle (Supplementary Fig. 1h).

### scRNA-seq profiling of *Cdh5*-CreERT^2^ marked VE-cadherin (*Cdh5*)^+^ expressing lineages

To understand the cellular heterogeneity and the dynamic molecular reorganization of VE-cadherin^+^ cells in hair cycle, we performed 10x Genomics scRNA-seq analysis of *Cdh5*-CreERT^2^ x tdTomato^+^; *Krt14-H2BGFP-* cells purified from mouse dorsal skin at quiescence (telogen) and growth (anagen) stages (Fig. 1c-e and Supplementary Fig. 1a), using methods previously described^29^. Two mouse replicates (S1 and S2) from each stage were subjected to 10x scRNA-seq library construction and sequencing (see Methods). After applying quality control parameters on individual samples (Supplementary Fig. 1i), we obtained a total of 7250 and 9262 high-quality cells at telogen and anagen respectively. All samples were integrated using the *Harmony* package (see Methods), followed by Uniform Manifold Approximation and Projection (UMAP) analysis which produced 16 distinct cell populations from the tdTomato^+^/GFP^-^ cells, highly reproducible in all 4 mice analyzed (Fig. 1f and Supplementary Fig. 1j). Of the 16 populations, 6 were *Cdh5* negative contaminating immune cells and fibroblasts, 3 populations were non-endothelial *Cdh5-low expressing cells* (Fig. 1h and Supplementary Table 1), while 7 were identified as bona-fide endothelial cells based on robust Cdh5 expression and other lineage markers (Fig. 1g,h). Previously characterized EC type-specific markers revealed distinct 30, 31, 32 populations of artery, vein, capillaries, and lymphatics^30,31,32^ (Fig. 1h). The capillaries (Cap) displayed two (*Cxcl12^+^/CD36^+^*)) populations enriched in either Vegfr2 (*Kdr*) or *Rgcc*^9,30^. The lymphatics were either collecting vessels (Lymph-Col, *Prox1*^+^) or capillaries (Lymph-Cap, *Ccl21a, Pdpn* and *Lyve1*^+^). The Ki67^+^ proliferative (Proli) EC population was ~1-2% of total, which is in good agreement with our BrdU data (Fig. 1f, g and Supplementary Fig. 1g).

After we computationally removed the non-endothelial populations, the 7 ECs populations represented 2543 (telogen) and 7498 (anagen) high-quality sequenced cells (Supplementary Fig. 2a) with high correlation between replicates (Supplementary Fig. 2b). Given previous analysis that lacked skin suggested tissue-specific expression of ECs^9^, we examined individual population enriched genes and compared them with other tissues. We found high enrichment of *Mgp* and *Stmn2* for Artery, *Aqp1* and *Sele* for Vein, *Gpihbp1* and *Rbp7* for Cap-Kdr, *Mmrn1* and *Lcn2* for Lymph-Cap, *Fibin* and *Cldn11* for Lymph-Col and many others (Supplementary Fig. 2c, d and Supplementary Table 2), and they significantly deviate in their expression patterns from ECs in other tissues. These markers may help in the future to elucidate skin-specific roles of these markers in ECs during skin homeostasis. Together, these data provide high-quality single-cell transcriptomic profiles for adult VE-cadherin^+^ skin lineages at different hair cycle stages.

### Adult skin endothelial populations undergo dynamic redistribution and gene expression changes during hair cycle

To probe in-depth the cellular and molecular heterogeneity underlying vascular ECs remodeling in hair cycle we first examined the distribution of different cell types at telogen and anagen. The UMAP analysis showed that all 7 EC populations that include the proliferative cells are present at both time points (Fig. 2a), with no emergence or loss of any EC type. Interestingly, the relative fraction of each population changed at anagen when compared to telogen, with a near doubling of blood EC populations at the expense of the lymphatic populations (Fig. 2b). To test for this change *in vivo*, we co-stained skin tissue at telogen and anagen with CD31 and either Endomucin (blood vessel - BV- marker) or Prox1 (lymphatic vessel - LV-marker). Using 8μm thin sections to reduce overlap, we identified nuclei enclosed by CD31 signal and counted BV and LV EC density in both dermis and hypodermis (Supplementary Fig. 2e, f). The data showed significant increase in BV EC density by ~45% at anagen, mainly in the hypodermis area (Fig. 2c, d). On the contrary, the LV EC counts showed decreased density at anagen mostly detected in the dermis (Fig. 2c, d). This *in vivo* validation by IF counts confirmed the population fractional change between the hair cycle rest and growth phases predicted by the scRNA-seq data. The reduction of lymphatics density at anagen is intriguing and may be related to the reported reduced drainage function at anagen^25, 26^. The increased blood vessel density at anagen aligns with proposed high energy needs to supply O2 and nutrients for the production of new hair shafts^22^ and to expand the skin hypodermis.

**Fig. 2.**
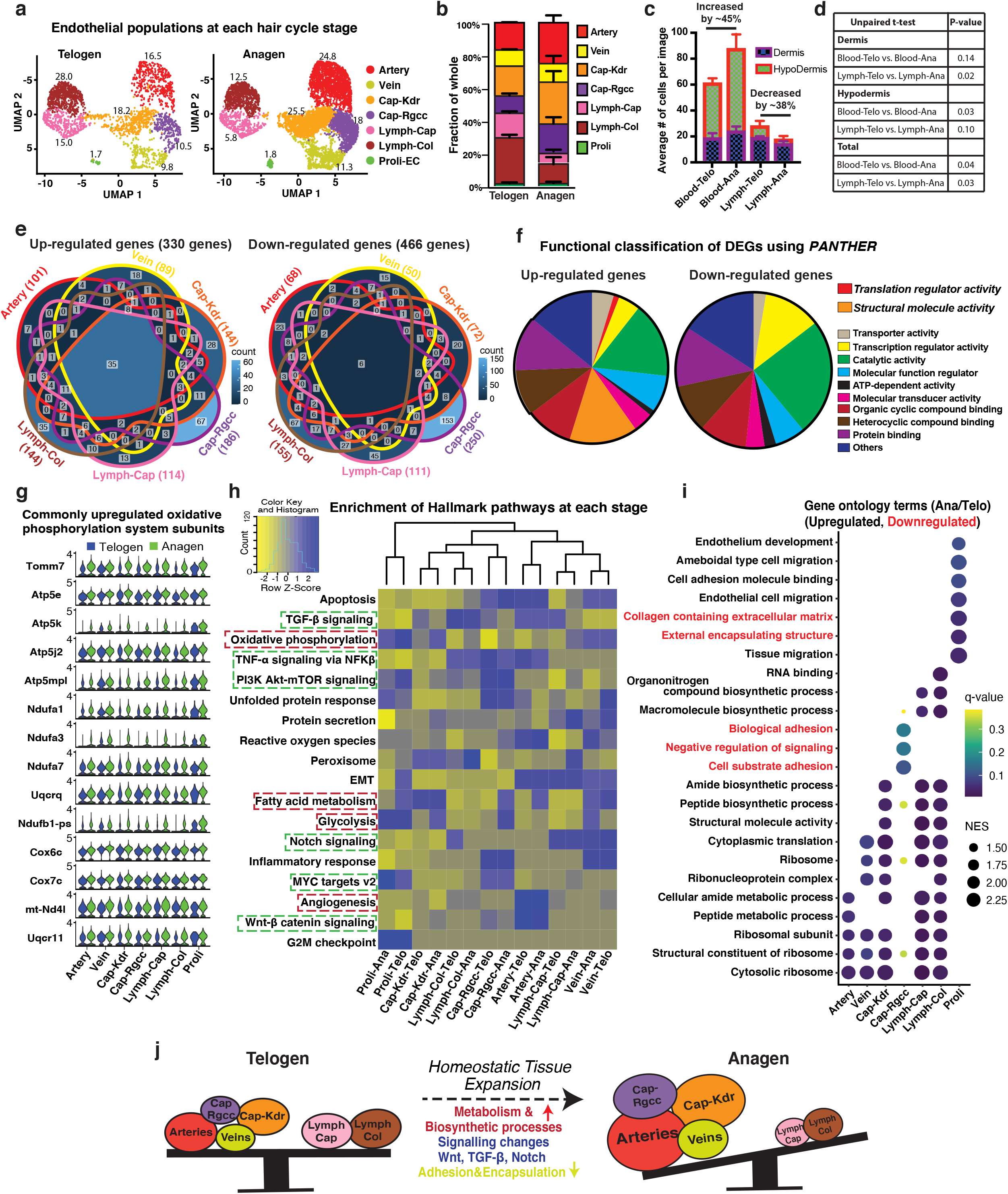
Dynamic changes of gene expression in endothelial populations at telogen to anagen transition. **a** UMAP plot showing 7 endothelial populations detected at telogen and anagen stages. **b** Fraction of each population (from total 100%) in scRNA-seq datasets showing increase of blood vessel proportion at anagen. **c** Quantification of blood and lymphatic EC numbers shows density per defined tissue length (per image). Data was split in dermis and hypodermis regions and was obtained by immunofluorescence staining for CD31 and either Endomucin (blood) or Prox1 (lymphatics) of thin (8μm) skin sections. Unpaired student t-test was used to directly compare each compartment between the two stages (N~30 images from n=3 mice/stage). **d** Table showing p-value significance for graph in **(c)**. **e** Venn diagram for differentially expressed genes (DEGs) that changed expression at telogen-anagen transition (FC>1.3) in non-proliferating endothelial populations. **f** Pie chart showing functional categories of up-and down-regulated genes using the *PANTHER* classification system. **g** Differential expression of oxidative phosphorylation system components at telogen and anagen. **h** Hallmark pathway analysis using all detected genes (raw counts obtained using *FetchData* function in *Seurat*) at each stage. The enrichment score of all hallmark pathways in all populations was compared by heatmap2 function in R. **i** Dotplot compilation of Gene Ontology (GO) terms analysis results performed using DEGs (FC>1.3) at anagen. Red fonts indicate downregulated categories. **j** Cartoon summarizing cellular and functional changes in endothelial populations while they transit from telogen to anagen stage *in vivo*.

To extract molecular signatures of individual EC populations and identify pathways that may change in hair cycle, we used a cutoff of log2FC>0.25 and noted substantial gene changes in each population at the telogen-anagen transition (Supplementary Fig. 3 a). Next, we extracted differentially expressed genes (DEGs) (330 upregulated and 466 downregulated or 784 total unique genes >1.3x) from non-proliferative populations (Fig. 2e and Supplementary Table 3). Classification of these DEGs using *PANTHER^33^* revealed diverse functional categories such as transcription, catalytic activity, and protein binding common to both upregulated and downregulated genes, while translation regulators and structural molecules activities were only in the anagen upregulated genes (Fig. 2f). Heatmap clustering of these DEGs (1.3x) illustrates the patterns of gene expression changes from telogen to anagen in the ECs (Supplementary Fig. 3b). Not surprisingly, the LV and BV segregate away from each other, and the Lymph-Cap and Lymph-Col vessels DEGs follow a similar pattern from telogen to anagen. Interestingly, the Cap-Kdr population clustered with the proliferative cells and are more closely associated with the arteries. Conversely, the Cap-Rgcc population clustered with the veins and away from the proliferative cells (Supplementary Fig. 3b).

A small fraction (35 genes) involved in metabolism and oxidative phosphorylation were upregulated in all EC populations at anagen, indicating increased cellular bioenergetics (Fig. 2g and Supplementary Table 3). Furthermore, *Hallmark* pathways^34, 35^ analysis of all detected genes show EC-population specific enrichment, and some change between anagen and telogen (Fig. 2h, Supplementary Fig. 3c and Supplementary Table 4). Interestingly, the artery population showed strong enrichment for angiogenesis, Wnt, TGF-β, and TNF-α pathway (Fig. 2h). Pseudotime analysis by Monocle 2^36^ also predicted the artery population as the ‘ground state’ or the most immature on the lineage trajectory (Supplementary Fig. 3d). Multiple signaling pathways such as TGF-β, NFK-ß, PI3K-Akt-mTOR, Notch and MYC pathways were also changed in different EC population (Fig. 2h). Finally, gene ontology (GO) terms analysis of genes upregulated (FC>1.3) showed anagen-enrichment for various biosynthetic processes in most populations but not in Cap-Rgcc, which displayed down-regulation of adhesive functions (Fig. 2i). Similarly, proliferating ECs showed enrichment for the endothelial development as well as cell migration and down-regulation of cell adhesion, extracellular matrix, structural encapsulating functions, as may be expected of vasculature growth at anagen (Fig. 2i). Interestingly, recognized angiogenesis-related genes^37^ were most enriched in the artery population, and did not change much at the two hair cycle stages (Supplementary Fig. 4). This result, along with the BrdU incorporation at telogen/early anagen (Supplementary Fig. 1g), and comparable percent of Ki67^+^ ECs at telogen and anagen (Fig. 2b) suggest that angiogenesis likely occurs at both stages, not just at anagen. Together, these analyses demonstrate significant changes in cellular fractions and absolute density of blood and lymphatic ECs, as well as in gene expression for multiple pathways that occur in vasculature during normal skin homeostasis (Fig. 2j). These data provide the molecular platform for probing regulation of adult vasculature control during tissue homeostasis and has implications to other highly regenerative systems besides skin (see Discussion).

## Discussion

Our work strengthens the current understanding of the adult skin vasculature system. This work should aid in efforts to engineer proper organotypic skin cultures for chronic and diabetic wounds^38^ and for understanding skin vascular diseases such as Human hereditary hemorrhagic telangiectasia (HHT)^39, 40^. Our data demonstrate that in homeostasis skin EC populations are not only remodeled, but they also coordinately change their molecular makeup to reflect increased metabolic and biosynthetic activities. This corroborates with previous findings of microvasculature remodeling through capillary pruning and duplication^41^ and through exercise-induced angiogenic stimulus^42^. Several major signaling pathways surfaced from our transcriptomic analysis as differentially regulated in the specific EC populations, including TGF-β, Notch and Wnt signaling. These data constitute a rich base for future investigation of the mechanisms regulating skin vasculature remodeling and the emerging cross-communication of ECs with hair follicle stem cells^15^, which are poorly understood. Interestingly, we detected an anagen-specific increase in blood vasculature at the expense of the lymphatic vasculature, suggesting the two lineages are in opposite flux during skin homeostasis. While the increase of blood vessels as hair grows is not surprising^15^, the concomitant decrease in lymphatics is intriguing. A reported reduction in the dermal lymphatic capillaries area at anagen^26^, and the reduced drainage observed at anagen^25^ corroborate our finding. The lymphatic vasculature is generally static in adult tissues^43, 44^, but homeostatic regeneration also occurs in the intestinal ‘lacteal’ lymphatic capillaries and the ovary during folliculogenesis^45, 46^. Our study in skin adds important insight into active vascular homeostasis of another highly regenerative adult tissue, suggesting adult vasculogenesis may be a broader process than previously realized.

## Methods

### Mice and treatments

All the mouse work was performed in accordance with the Cornell University Institutional Animal Care and Use Committee (IACUC) guidelines (protocol no. 2007-0125). The tdTomato^28^ mice were obtained from Jackson Laboratory. The *Cdh5*-CreERT^2^^27^ mice were imported from Dr. Anne Eichmann at Yale University and used as previously described^20^. The mice were intraperitoneally (IP) injected with tamoxifen at dose of 200μg/g body weight to induce Cre recombination. For BrdU-pulse experiments, mice were fed with 0.8mg/ml BrdU in daily water supply.

### Immunofluorescence staining, microscopy and image processing

Immunofluorescence (IF) staining on the non-prefixed tissue sections was performed by following the standard protocol, as described previously^20^. The tdTomato^+^ skin tissues were prefixed in 4% paraformaldehyde for 2 hours at 4□ followed by passing them through a sucrose gradient before embedding in OCT compound (Tissue Tek, Sakura). The non-prefixed samples were first fixed in 4% paraformaldehyde (PFA) for 10 minutes at room temperature (RT), followed by washing with PBS and 20mM glycine in PBS. The sections were blocked with 5% normal serum for 1 hour at RT before incubating them with primary antibodies at 4□ overnight. Next day, sections were 3x washed with PBST and then incubated with secondary antibodies (1:500). The sections were washed 3x with PBST and stained with DAPI. The slides were mounted with antifade solution. For thick sections (70μm), primary antibodies incubation was performed for 48 hours at 4□ and secondary antibodies overnight at 4□. For BrdU staining, tissue sections or cells were first stained with non-BrdU primary antibodies, followed by fixation using 4% PFA. Chromatins were denatured by incubating the slides in 1M HCl solution for 55 minutes at 37□. The slides were washed with PBS and re-blocked using 0.5% Tween-20 and 1% BSA before incubating them with anti-BrdU antibody overnight at 4□. Then slides were washed in PBST and incubated with FITC-conjugated anti-rat antibody. Primary antibodies used in this study include CD31 (1:100, BD Biosciences, 550274); VE-cadherin (1:500, R&D Systems, AF1002); LYVE1 (1:400, Thermo Scientific, 14-0443-82); Prox1 (1:300, Abcam, ab199359); BrdU (1:300, Abcam, ab6326); Endomucin (1:300, Santa Cruz Biotechnology, sc-65495). The IF images were captured using either the Leica DMI6000B microscope with Leica K5 camera or the Zeiss LSM inverted 880 confocal microscopes. The z-stacks (1μm stacks) images of thick sections were deconvoluted using AutoQuant X software and were further processed for brightness, contrast and stitching using FIJI^47^. Multiple images were combined using stitching package^48^ in Fiji^47^ for Fig. 1. The quantification graphs were generated using GraphPad Prism 9.

### Fluorescence-activated cell sorting (FACS) of VE-cadherin^+^ cells

FACS isolation of VE-cadherin^+^ cells and its purity check was performed as previously described^29^. Briefly, tdTomato (Jax Stock #007905), *Cdh5*-CreERT^2^^27^ and *Krt14* -H2BGFP^49^ mice were used for VE-cadherin^+^ cells and control keratinocytes isolation. The VE-cadherin^+^ cells were labeled with tdTomato by injecting tamoxifen (200 μg/g) at postnatal day (PD)17, and dorsal skin was collected at PD20 or PD32. The dorsal skin was digested in collagenase and Dispase mixture as previously described^29^. The dead cells were removed by LIVE/DEAD™ Fixable Aqua Dead Cell Stain Kit (ThermoFisher). FACS Aria (BD Biosciences) was used for the cell sorting in Cornell Flow Cytometry facility. FACS data were analyzed with the FlowJo (FlowJo™ Software, v10.5.0, BD Biosciences).

### Single Cell library preparation and data analysis

The barcoded single-cell 3’ cDNA libraries were generated using Chromium Single Cell 3’ gel bead and library Kit v3 (10x Genomics) and sequenced using an Illumina NextSeq-500. The raw data were aligned to the mouse reference genome (mm10-2020-A) using the 10X Genomics *Cell Ranger* pipeline (v6.0.1). The data analysis was performed in R using the Seurat package version 4.0^50^. Cells that had between 200 and 5000 genes expressed and had under 10% of the UMIs mapped to mitochondrial genes were retained. We obtained a total of 3957 and 8627 high-quality cells for telogen and anagen samples, respectively. For further analysis, all samples were merged, the transcript counts were log-normalized, and the expression of each gene was scaled so that the variance in gene expression across cells was one, followed by Principal Component Analysis (PCA). The data integration was performed by PCA embeddings using *Harmony^51^*. The corrplot R function was used to measure the correlation between the two biological replicates. Differentially expressed genes (DEGs) were identified by *‘FindAllMarkers’* function using the Wilcox Rank Sum test.

### Cell lineage trajectory, gene scoring and gene ontology analysis

Trajectory analysis was performed using Monocle2 and Monocle3^36, 52^. We used the top 2000 genes to order cells in pseudotime trajectory by Monocle2 and the cells having lower than 200 transcripts were removed from further analysis. The DDRTree package was employed to construct the trajectories. To determine the collective enrichment of a set of genes in different populations, the AddModuleScore function in the Seurat R package was used. For pathway analysis, DEGs were identified by Seurat’s ‘FindMarkers’ function using the Wilcox Rank Sum test (FC>1.3). Further enrichment analysis and plotting were performed as previously described^29^.

### Heatmap of Differentially expressed genes (DEGs) using raw counts

First, DEGs were identified using *FindMarkers* function of *Seurat*. Next, to obtain the expression value for each gene in all populations, the raw counts for the DEGs were extracted using the *FetchData* function. Of 205 genes (FC >1.5) 6 showed 0 values in at least one population, whereas the corresponding population in the other stage showed a very low expression level relative to other populations. Thus ‘0’ was replaced with the low ‘background’ value from the corresponding population at the other stage. This way, these genes with very low expression levels in some populations appear ‘unchanged’ from telogen to anagen, thus eliminating the possibility that small fluctuations in background expression are assigned a ‘changed’ value. The *Xist* gene showed 0 values in more than one population but did not display an obvious background value in the corresponding population at the other stage, and thus they were not analyzed further. The same method was applied to the gene set of 784 genes (FC>1.3), where 18 showed 0 values in at least one population.

## Data availability

The scRNA-seq data have been deposited in the NCBI Gene Expression Omnibus (GEO) database with accession numbers GSE211381. The records are accessible at: https://www.ncbi.nlm.nih.gov/geo/query/acc.cgi?acc=GSE211381

## Code availability

This paper does not report original code. Any additional information required to reanalyze the data reported in this paper is available from the lead contact upon request.

## Acknowledgments

We thank Lauren D. Walter and Benjamin D. Cosgrove from Cosgrove laboratory for their help with scRNA-seq analysis and helpful suggestions. We also thank the Center for Animal Resources and Education (CARE) for mouse care and the Cornell University Biotechnology Resource Center (BRC) for their help in scRNA-seq experiments. Confocal microscopy work on Zeiss upright 880 was supported by the Cornell BRC facility (NYSTEM C029155 and NIH S10OD018516). This work was supported by NIH/NIAMS Grants R01AR070157, R01AR073806 and R56AR081021 to TT and by the Empire State Stem Cell Fund through New York State Department of Health Contract # C30293GG training grant to GC. Opinions expressed here are solely those of the author and do not necessarily reflect those of the Empire State Stem Cell Board, the New York State Department of Health, or the State of New York. We thank Jonathan Li for his help in immunofluorescence stainings and for his assistance in manuscript editing.

## Author contributions

GC, NL and TT designed the experiments. GC and NL performed the experiments. GC did the computational analysis and *in vivo* validations. SG helped with pathway analysis. GC and TT analyzed and interpreted all data. GC and TT prepared the figures. GC and TT wrote the manuscript.

## Supplemental Figures legends

**Supplementary Fig. 1, related to Fig. 1.**
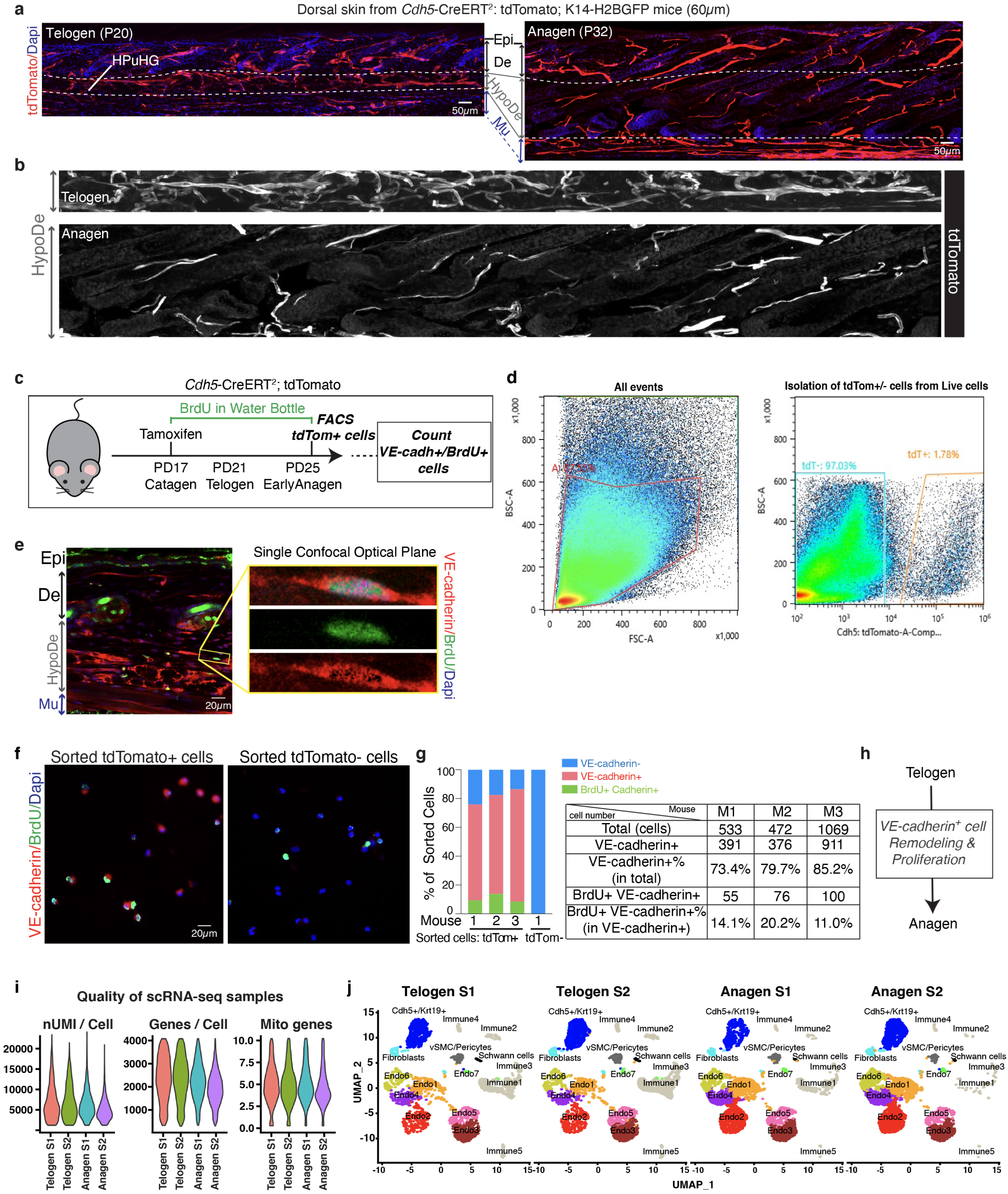
Proliferation of VE-cadherin^+^ cells *in vivo* and their FACS isolation for scRNA-seq analysis. **a** Maximal projections of confocal stack images showing 60μm dorsal skin sections at telogen and anagen from *Cdh5-*CreERT^2^; tdTomato (red); *Krt14-H2BGFP* mice (green channel not shown). HPuHG area is circled in a white dotted line. Scale bar 50μm. HypoDe, hypodermis; Mu, muscle; De, dermis, Epi, epidermis. **b** Single channel image (tdTomato) of cropped HPuHG region from **(a)**. Note dispersion of HPuHG in the hypodermis area (HypoDe) area. **c** Schematic for tamoxifen induction and sample collection for BrdU-pulse experiment. **d** Gating strategy for isolation of tdTomato^+^ cells from *Cdh5*-CreERT^2^; tdTomato mice for BrdU incorporation assay *in vivo*. **e** Confocal single optical plane of 30μm thick skin sections from BrdU-pulsed mice stained for VE-cadherin (red), BrdU (green) and Dapi (blue). Enlarged cropped images on the right show BrdU in VE-cadherin^+^ cell. **f** Sorted tdTomato-positive and tdTomato-negative cells stained with VE-cadherin (red), BrdU (green) and Dapi (blue). **g** Quantification for VE-cadherin and BrdU stainings co-localization in sorted cells from panel **(f)**. n=3 mice. **h** Summary showing tdTomato^+^ cells undergo remodeling and proliferation from telogen to anagen transition *in vivo*. **i** Quality of selected cells from two mouse replicates per stage that were used for the scRNA-seq analysis, nUMI/Cell – Unique Molecular Identifiers detected per cell, Genes/Cell - number of genes detected per cell, and Mito genes – a percentage of mitochondrial genes per cell, S – sample. **j** First-level UMAP clustering identified a total of 16 populations in both replicates S1 and S2 at each stage (the combined UMAP was split by sample).

**Supplementary Fig. 2, related to Fig. 1. and 2.**
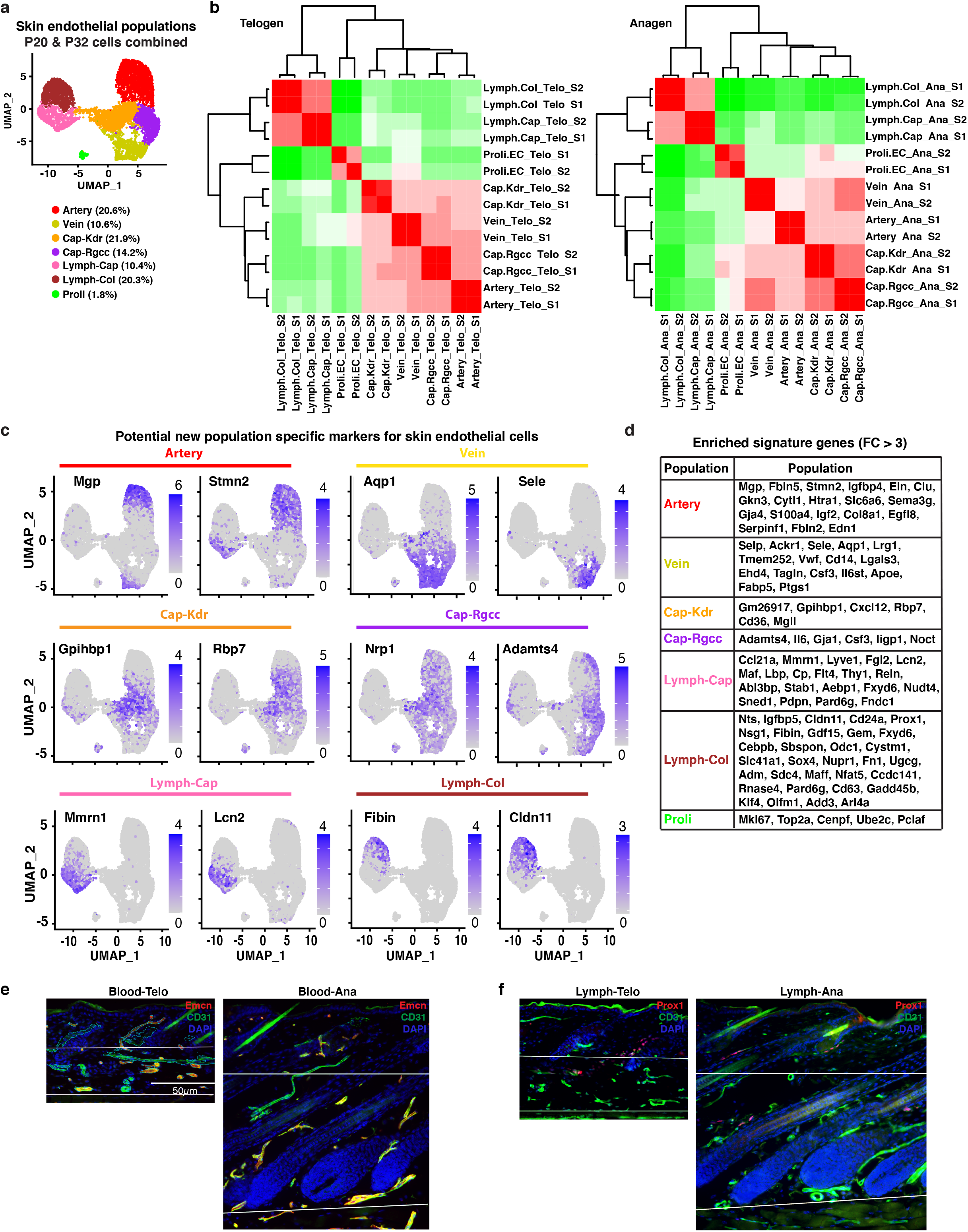
Decoding identity of single cell populations and counting of blood and lymphatic ECs *in vivo*. **a** UMAP plot showing 7 endothelial populations after computationally removing the non-ECs followed by re-clustering (all four samples combined before generating UMAP plot). **b** Comparison of gene expression matrix from two replicates for all 7 endothelial populations that showed a high correlation between samples at each stage. **c** Feature plots showing expression and distribution of selected signature genes for each endothelial population. **d** Table for top upregulated signature genes for each population. (FC>3). **e-f** Immunofluorescence staining images of 8μm thin skin section from PD20 (telogen) and PD32 (anagen) mice stained with either Endomucin (EMCN, red), CD31 (green) and Dapi (blue) (**e**) or Prox1 (red), CD31 (green) and Dapi (blue) (**f**). The whole skin was divided into dermis and hypodermis by a white line and the stained structures were encircled for nuclei counting purpose, as represented in Blood-Telo image. Scale bar 50μm.

**Supplementary Fig. 3, related to Fig. 2.**
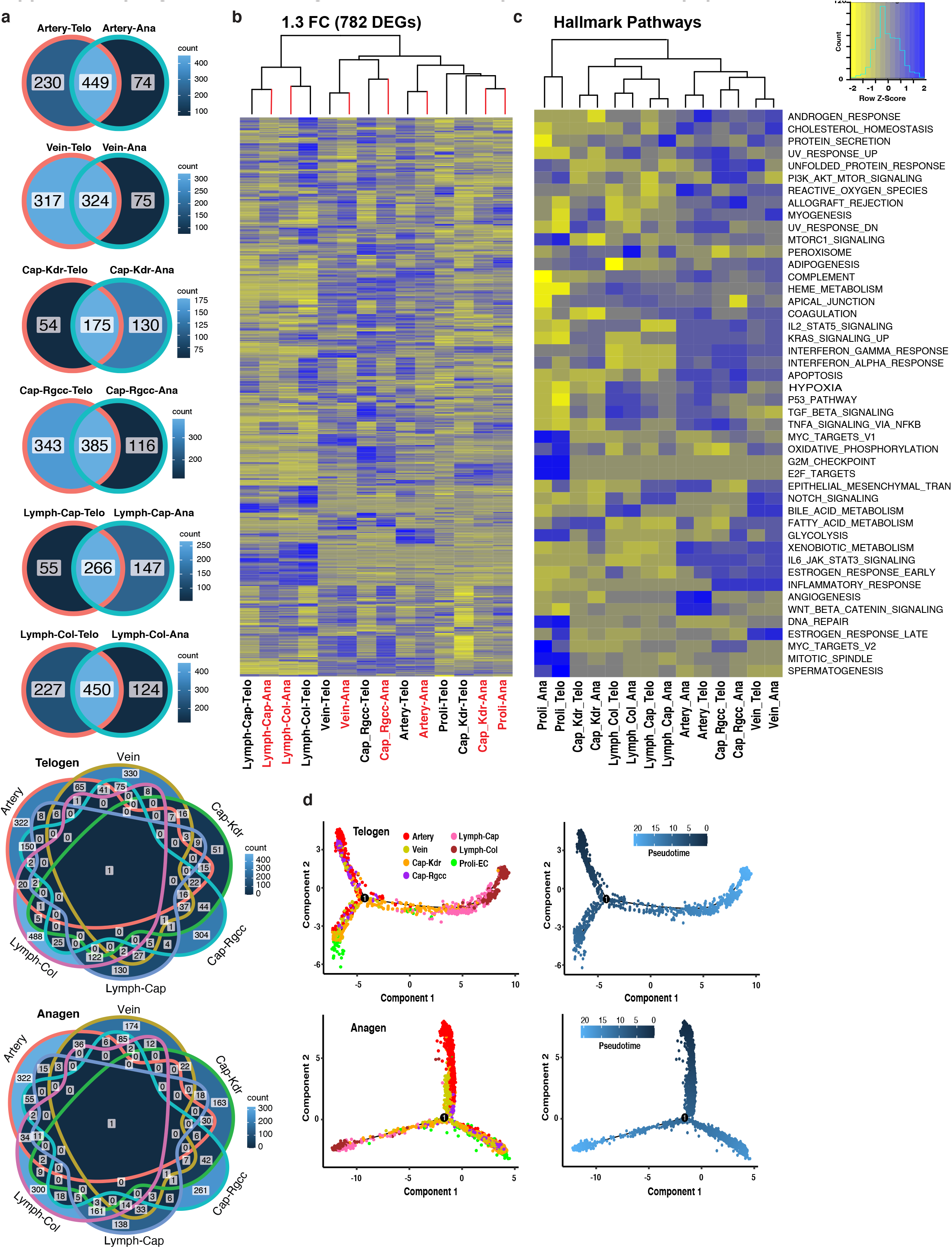
Functional analysis on endothelial populations. **a** Venn diagrams for signature genes (log2FC>0.25) at indicated stages. The signature genes were obtained by comparing each population with the rest of the populations at the same stage. The first six plots show individual populations at two stages; last two plots show all populations at each stage. **b** Heatmap clustering of 782 DEGs (FC>1.3) showing population clustering and extent of gene expression changes between two stages (Raw values of DEGs in each population were extracted using *FetchData* function). **c** Raw hallmark pathway analysis using all detected genes at both stages. The enrichment score was compared using the heatmap2 function in R. **d** Single-cell transcriptome trajectory reconstruction of EC populations using Monocle2 to predict lineage transition path. Pseudotime coloring indicates ground state in dark blue, corresponding to artery.

**Supplementary Fig. 4, related to Fig. 2.**
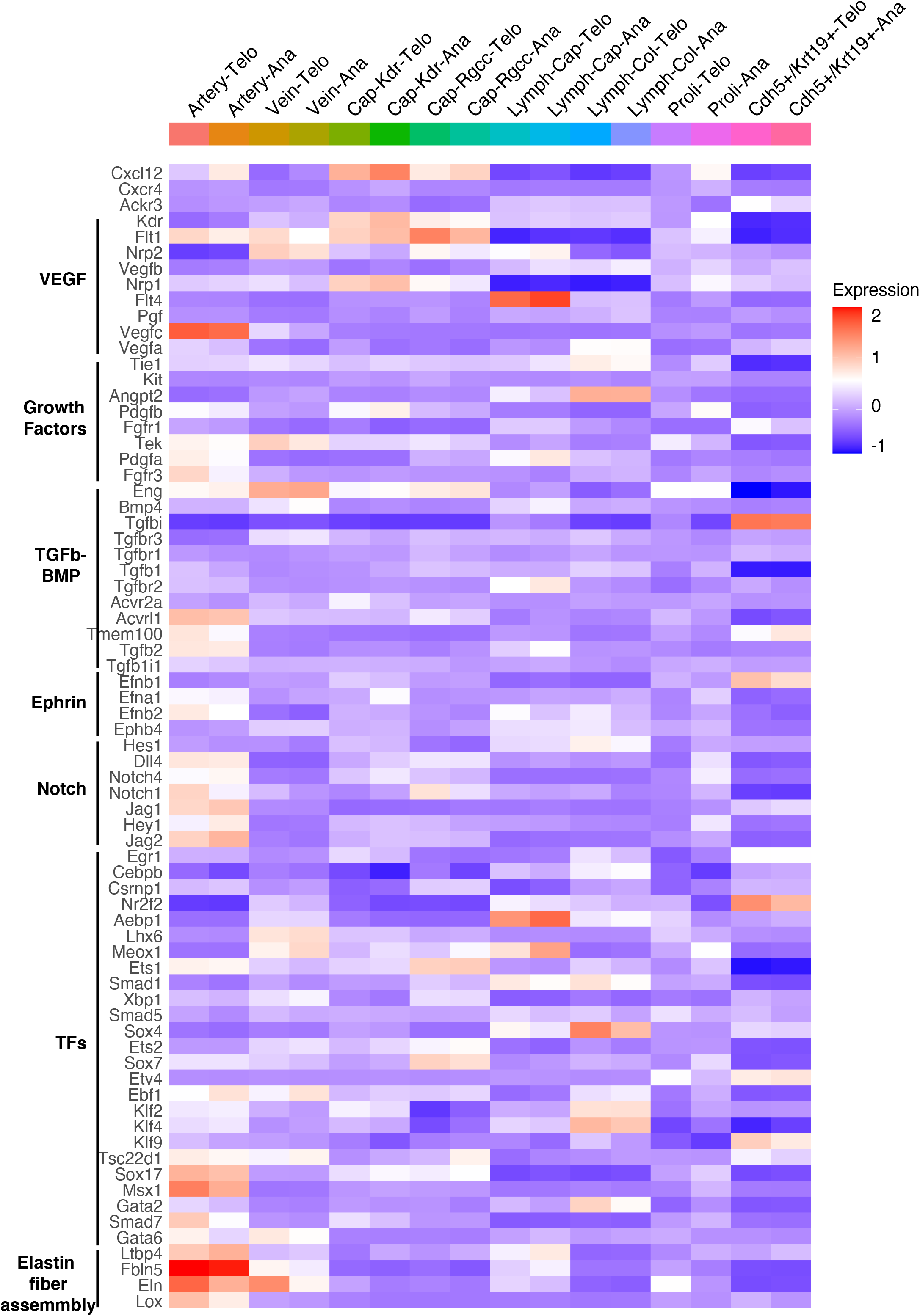
Expression of angiogenesis-related factors in skin EC populations. Heatmap showing expression of various gene classes known to be involved in angiogenesis. The gene list was extracted from Brulois *et al*., 2020.

